# Texture-Based Image Analysis For the Assessment of Soybeans

**DOI:** 10.1101/2023.01.20.524975

**Authors:** Alexander Lee, Kevin Hoffseth

**Affiliations:** Department of Biological and Agricultural Engineering, Louisiana State University AgCenter, 149 E.B. Doran Building, Baton Rouge LA, 70803, USA

## Abstract

At 4.44 billion bushels worth an estimated $57.7 billion dollars, soybeans are one of the most produced crops in the United States and are appraised using a standard grading system set by the USDA(NASS, 2022). The grading and inspection process is based on twenty factors, including six that are visual based (USDA, 2020). The aim of this research is to apply texture-based image analysis to assess visual indicators of damage in soybeans and demonstrate potential use in increasing efficiency and consistency in soybean grading. Surface texture is one of the characteristics that is examined by inspectors as part of assessing soybean damage, while “image texture” is a calculated set of parameters that are used in image processing and analysis to quantify the apparent actual texture captured in an image. In this study, texture analysis using Haralick textural features is performed on sets of soybean images to assess damage types (as defined by the USDA inspection handbook and visual reference images) and improve soybean classification.

## Introduction

The United States produced 4.44 billion bushels of soybeans in 2021 worth an estimated $57.5 billion, an all-time production high and a five percent increase from the year before (NASS, 2022). In 2021 Louisiana produced 55.1 million bushels worth $694.5 million (NASS, 2021). Soybeans are currently inspected and appraised using a standard grading system set forth by the USDA. The resulting grade influences the price at which a quantity of soybeans can be sold. This grading system considers several factors such as heating, odor, damage, and insect infestation among others when giving a grade to soybeans being sold (USDA, 2020). USDA certified inspectors take samples for grading directly from the transport truck and grade them at the point of sale. Inspectors use either a hand or mechanical probe to take a sample of at least 125g. The sample is first inspected for foul odors and insect infestation. A portion of the original sample is then separated, and the moisture content is determined. The sample is subsequently tested for dockage where the amount of damaged soybeans and materials other than grains (MOGs) is recorded(USDA, 2020). In order to determine dockages during the grading process the USDA uses a set of Visual Reference Images (VRIs) that provide the inspectors with an image of the different types of damages that can occur in soybeans (USDA, 2016). The soybean samples are compared to the VRIs to determine type and amount of damage. This visual determination of dockages in conjunction with amount of MOGs, odor, heating, moisture, and other factors are the basis of the inspector’s grade determination.

There exists a need for increased accuracy and automation in the grading process, with opportunity arising in improved visual evaluation of soybeans. Methods of image processing and analysis (Gonzalez R. C., 2008), which apply mathematical operations and algorithms to extract data from digital images are a natural fit for improving visual analysis. Image processing and analysis techniques have been used previously to detect damage found in soybeans, for example in an automatic method of counting and separating damaged soybeans using size, shape, and color (Wattana, Siriluk, & Khotwit, 2018), along with efforts to identify soybeans with mildewed, insect damaged, skin-damaged, broken, and partly defective kernels using color, shape, and texture analysis (Liu et al., 2015). Color features in the RGB color space have been used to classify soybean seeds into sound, heat damaged, green frost damaged, and stink bug damaged categories with increased accuracy (Shatadal & Tan, 2003). Mold damage of soybeans was identified based on the variance in light reflectance between the mold damaged and non- mold damaged areas on the surface (Gunasekaran, Cooper, & Berlage, 1988). The hue, saturation and intensity (HSI) color space has been used in conjunction with morphological operators to detect and segment MOGs, split soybeans, and contaminated soybeans(Momin, Yamamoto, Miyamoto, Kondo, & Grift, 2017).Near-infrared hyperspectral imaging has been used to nondestructively determine soybean viability(Baek et al., 2019). Image processing and analysis techniques have also been used to detect fungal disease symptoms in the leaves, stems, and fruit of various groups such as fruits, vegetables, commercial crops, and cereals(Pujari, Yakkundimath, & Byadgi, 2015).

Texture analysis is the characterization of regions in an image by quantifying qualities described by terms such as coarse or smooth as a function of the spatial variation in pixel intensities. Multiple methods have been developed to achieve this, such as grey-level co-occurrence matrixes (GLCM), Laplace filters, and granulometric analysis (Kupidura, 2019). Haralick features, the method used in this study, are based on the grey-level co-occurrence matrix (GLCM), a statistical method of characterizing the spatial relationship of pixels in a greyscale image(Haralick, Shanmugam, & Dinstein, 1973). The GLCM is constructed by recording how often two grey-level pixels appear next to each other in an image. While some of the features identified by Haralick “relate to specific textural characteristics of the image such as homogeneity, contrast, and presence of organized structure”, other features only “characterize the complexity and nature of the grey tone transitions (Haralick et al., 1973).”The Haralick features were originally and still are used to classify aerial and satellite images into various land use categories. In recent years Haralick features have been applied to medical imaging to discriminate lung abnormalities (Zayed & Elnemr, 2015).

Previous studies using image analysis to evaluate the quality of soybeans share a commonality in that the researchers conducting these studies decide what criterion their systems will judge the damaged soybeans by(Liu et al., 2015; Wattana et al., 2018). A recent study uses comparisons between damaged soybeans and the average of values of normal soybeans to identify and remove damaged soybeans, but provides no definition of what makes a soybean normal (Liu et al., 2015). A different method considers a soybean as damaged if the number of white pixels in a binary image of a soybean is less than a set value (Wattana et al., 2018). Using this method would result in small and irregular, but otherwise undamaged soybeans being considered damaged. Healthy and normal soybeans are a variety of shapes and sizes and drawback to existing approaches in classification of soybeans are the subjective definition of healthy or normal soybeans. Two issues arise when using arbitrary definitions of healthy. The first is that the quality of the soybean is compared to the averages of what are presumed to be good soybeans. The second is that these systems tend to be binary systems. Meaning the soybean in question either matches the “good” soybean or it is considered damaged. In practicality many of the damages looked at by inspectors are measured on a scale of values. Soybeans with mold damage for example are only considered damaged if the “soybeans contain mildew on fifty percent or more of the seed coat in sufficient concentration to meet or exceed the minimum(USDA, 2020).”

A method for visually identifying and quantifying damage in soybeans would provide information for a more accurate quality assessment based on the actual damage found, and not a comparison to a “perfect” soybean. This study proposes a method to image and apply Haralick texture analysis to soybeans in order to evaluate their ability to distinguish damage types. The aim is not to determine if the soybean is “good” or “bad”, but to ascertain if certain damages are present and which Haralick textural features provide the best distinction between the categories.

## Materials and Methods

### Soybeans

The soybeans used in this study were comprised of grade 1-4 samples obtained from the Louisiana Farm Bureau. The soybeans harvested in June 2020 were graded by a certified USDA inspector. The samples were stored in plastic bags placed inside plastic bins at an average temperature and humidity of 70&F and 55 percent humidity respectively. A total of one hundred and fifty soybeans were chosen for this study and imaged between August and October 2020. The damage status of each soybean was characterized manually through use of the USDA inspection process and VRIs. In order to ensure an unbiased categorization, the one hundred and fifty soybeans were arbitrarily numbered and characterized by in a blind process by five volunteers, categorizing the soybeans as either smooth, cracked, wrinkled, or other. Representative example images of each soybean damage type can be seen in Figure 1.

**Figure 1:**
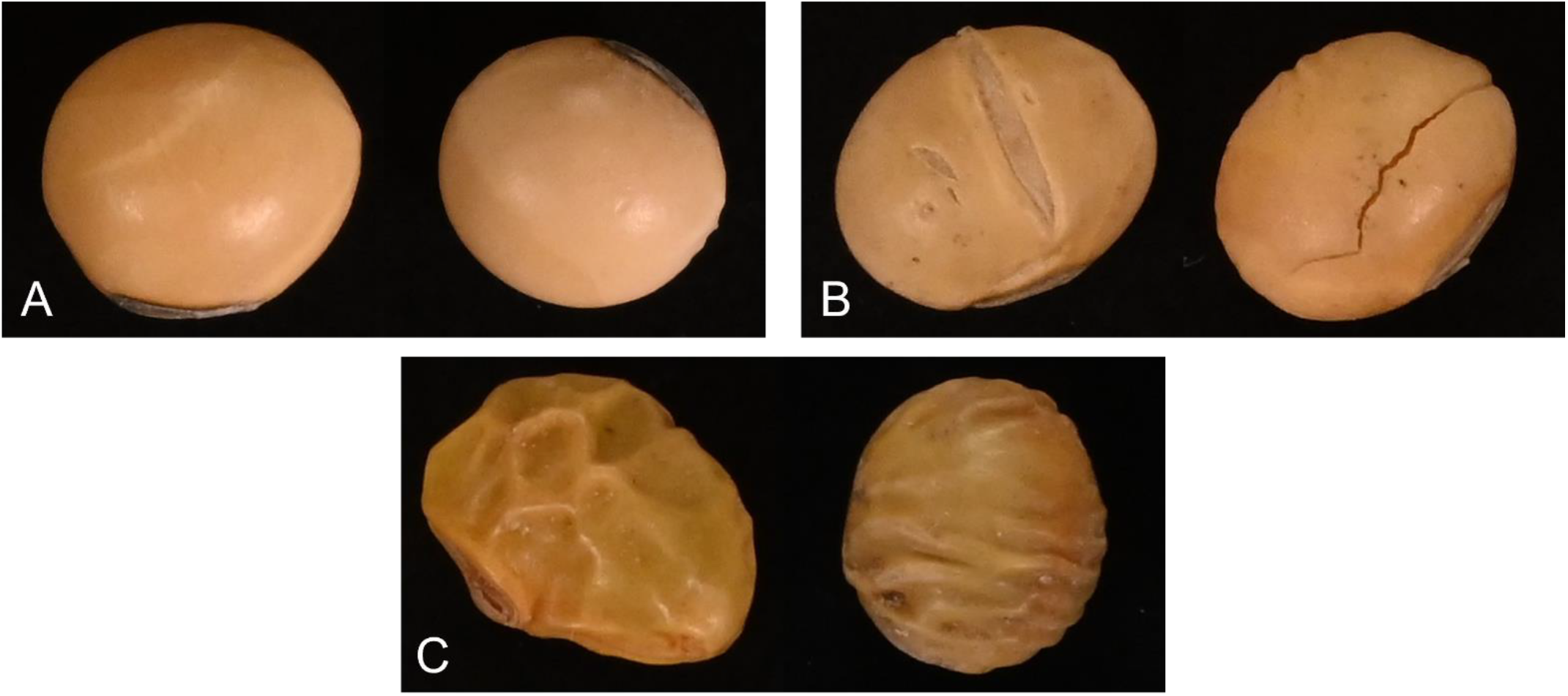
Examples of the categories used. Smooth (A), Cracked (B), Wrinkled (C)

### Image Acquisition

A 24.5-megapixel Nikon z6 camera with a Nikkor Z 24-70 mm lens was used to obtain images of the soybeans used in this study. All images were acquired with constant settings: a F-stop of 20, a shutter speed of 1/125, and an ISO of 2500. The camera was attached to a stand with the lens 21.4 cm above the imaging platform. The imaging platform consisted of a 36 cm by 30.5 cm plate of metal that was sanded smooth and painted matte black to reduce light reflection, shadows, and provide a high contrast to the soybeans. Three GVM-672 LED lights were used in the image acquisition setup, two behind the imaging platform angled at 45& and the third in directly in front of the platform. The temperature of the light was set to 3500 K& and the brightness was set to 50%. The actual measured temperature and brightness of the imaging platform was 3390 K& and 2890 lux respectively. The soybeans were placed in three rows of three, approximately 1-2 cm apart, allowing for nine soybeans per image. This provided reasonable imaging time versus single soybean resolution. The layout of the image acquisition setup can be seen in Figure 2.

**Figure 2:**
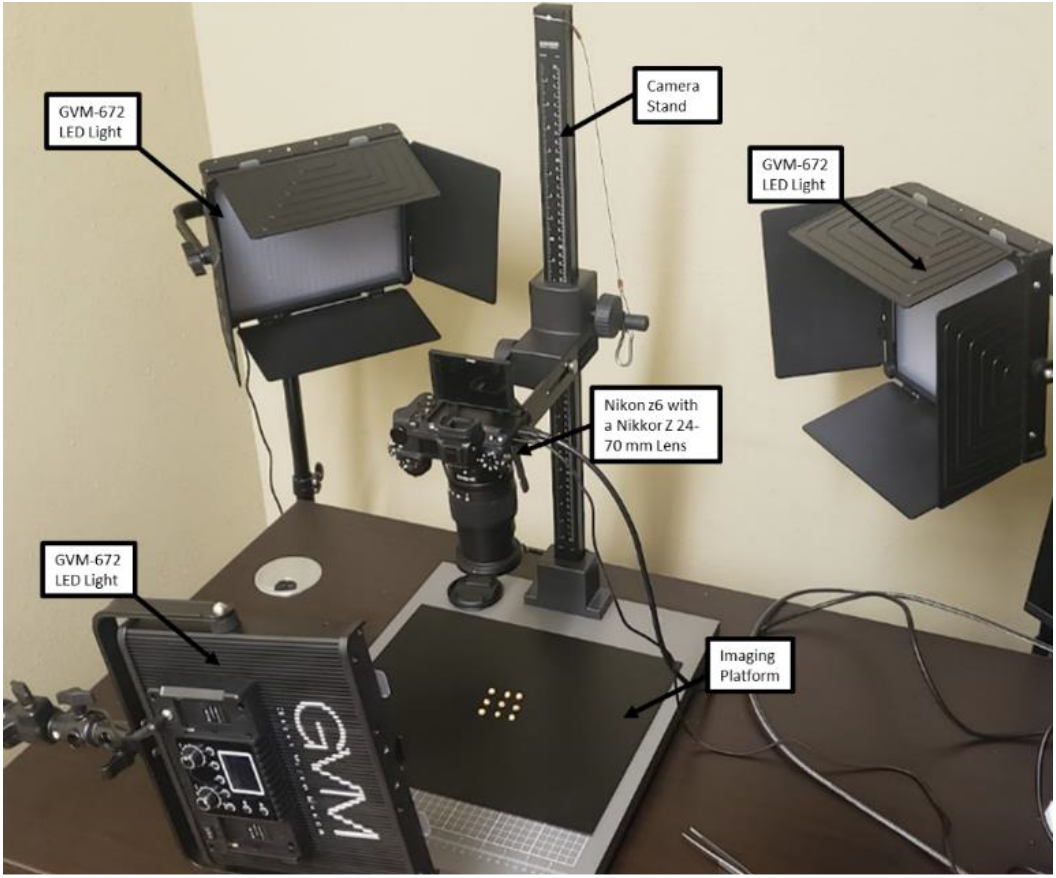
Layout of three light image acquisition setup

### Image Processing

After the images were acquired, they went through multiple processing steps. First the algorithm imports the original image and splits it into red, green, and blue channels. Through experimentation it was found that using only the red channel provided the most contrast, allowing for better detection of the edges of the soybean. Using the red channel, the edges were found using the Canny edge method(Canny, 1986). Then using the Canny edges the contours of each soybean are identified using the Suzuki method(Suzuki, 1985). A mask of the contours was placed over the original RGB image and a bounding box that encompasses each individual contour was applied. The bounding boxes were given a 140-pixel cushion and each box was segmented out of the original image. This created nine new images, each one containing a single soybean. This step was repeated for each of the original images. These color images are converted to greyscale using the weighted method. The weighted method weighs the red, green, and blue values of each pixel using the following equation: Grayscale = 0.299R + 0.587G + 0.114B (Bovik, 2010). A binary threshold is applied for Canny edge detection (Canny, 1986). The binary image then undergoes five iterations of both dilation and erosion using a two pixel by two-pixel kernel(Gonzalez R. C., 2008). Dilation and erosion iterations are used to remove artifacts created by other particulates in the original image such as shadows, dust, and dirt. After dilation and erosion, the resulting contours in the image are detected, counted, and the internal area is measured using the Suzuki method (Suzuki, 1985). The list of contours is then sorted based on the measured internal area and the largest contour, which must be the soybean, is selected. Using the calculated moment and centroid of the contour, an ellipse was fit to the contour using the algebraic distance algorithm (Fitzgibbon & Fisher, 1996). An inscribed rectangle was created using the computed major radius of the ellipse that was fit to the soybean contour. The inscribed rectangle is created for each of the soybeans and is segmented out for texture analysis. The step-by-step process is depicted in Figure 3.

**Figure 3:**
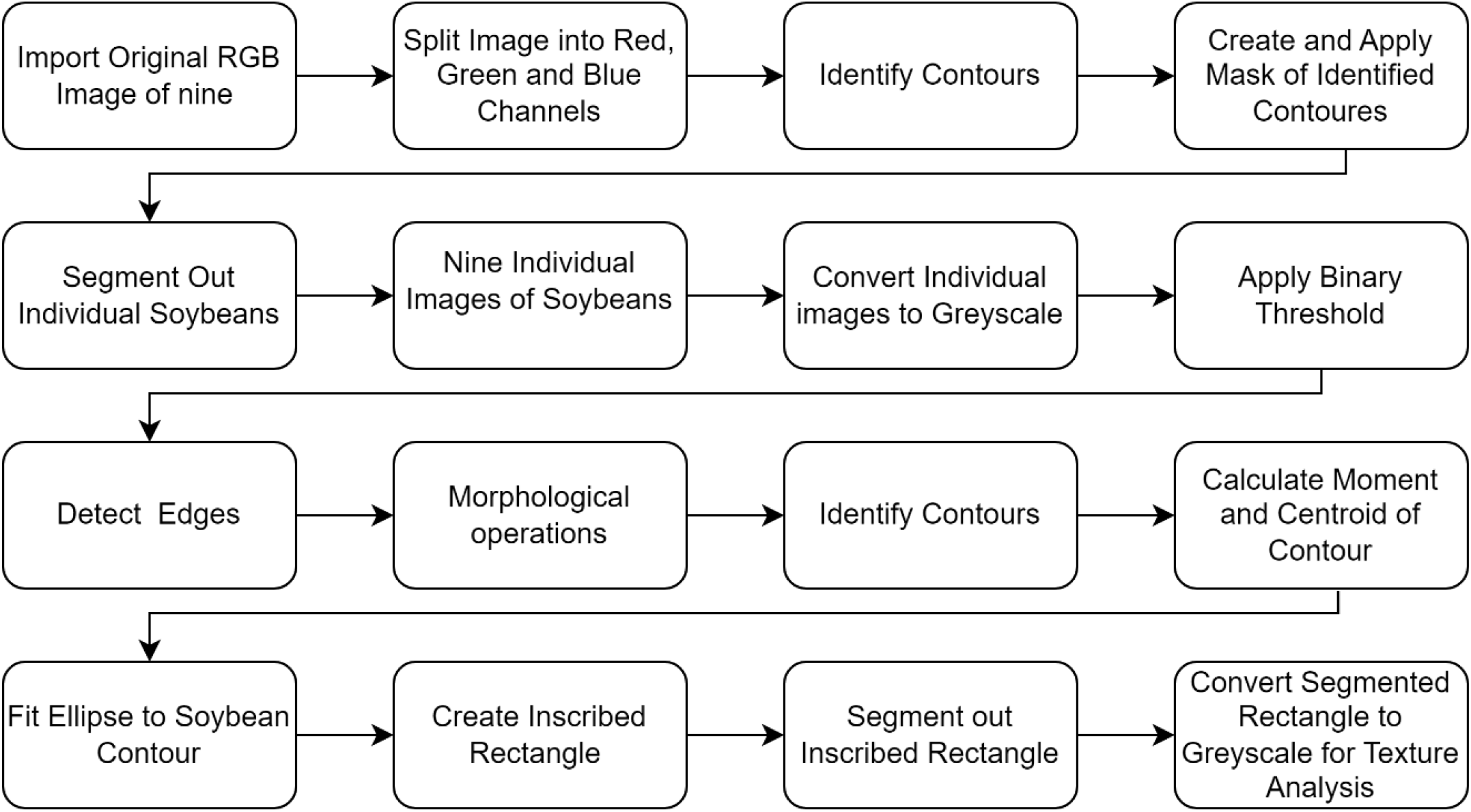
Flowchart of Image Processing Steps

**Figure 4:**
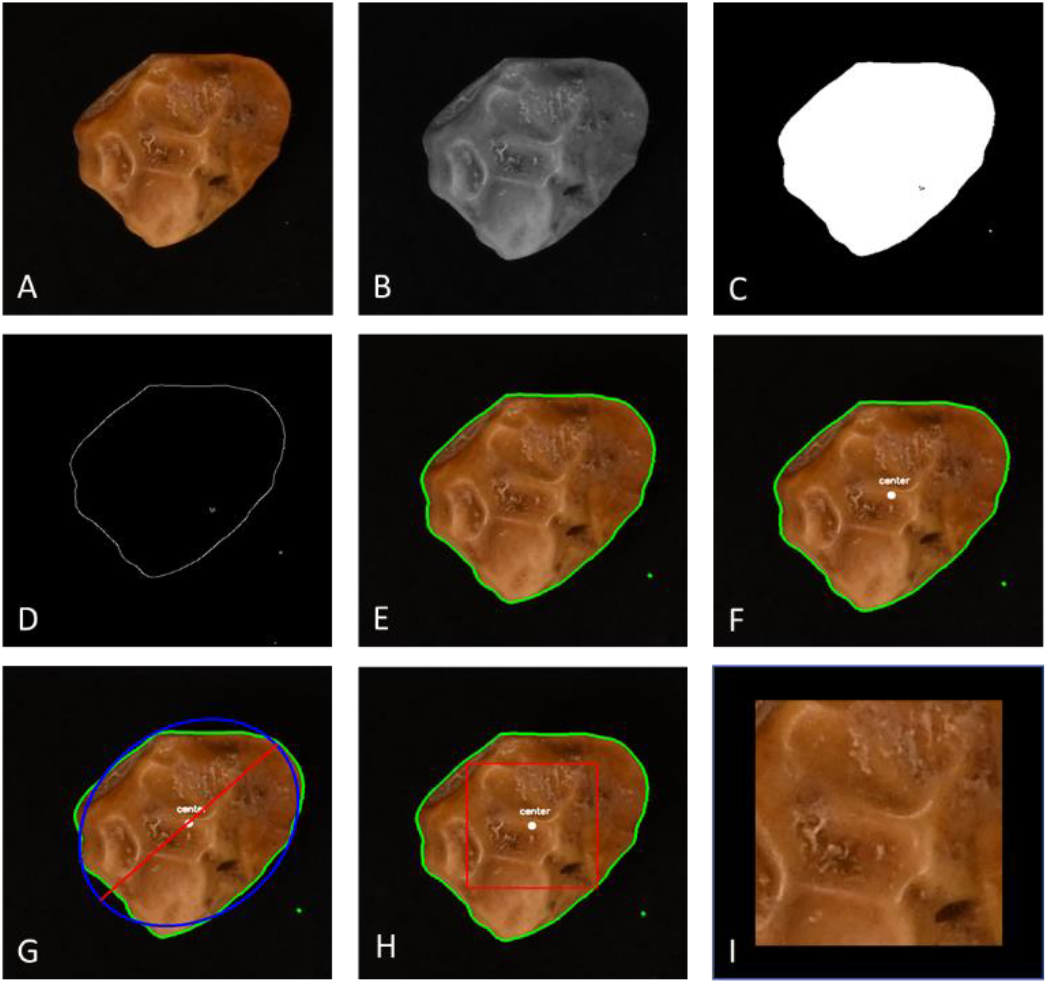
A- Original Image, B- Greyscale Image, C- Binary Threshold, D- Edge of the soybean, E- Contoured image, F- Contoured image with the calculated center, G- Soybean fit with an ellipse, H- Soybean with interior box, I- Segmented soybean

### Texture Analysis

The surface texture of the soybeans was examined using the 13 textural features detailed in Table 1, originally proposed by Haralick et al., in 1973. Haralick features rely on the notion that the properties texture and tone are related to each other and are both always present in an image. Tone is the level of brightness of each pixel ranging from black to white. Little variation in the features of an image indicates that the prevailing property is tone while larger variation would indicate that texture is the main property. Tone is ingrained the varying levels of greyscale pixels in an image, and texture is related to with the spatial or statistical distribution of the greyscale pixels (Haralick et al., 1973). Haralick textural features are a statistical approach to texture analysis that represents texture using non-predictive properties that control the relationships between greyscale pixels in an image(Materka & Strzelecki, 1998). The features are calculated by applying the equations for each feature, located in Table XX, to the grey-level co-occurrence matrix (GLCM). The GLCM is constructed by recording how often two grey-level pixels appear next to each other in an image. Using the greyscale center of each soybean that was segmented out at the end of the processing algorithm, the 13 textural features are calculated. The initial output is four values for each feature, one for each of the nearest neighbor pixels at 0, 90, 180, and 270 degrees. These four values are then averaged to get a single value for each Haralick feature.

**Table 1:**
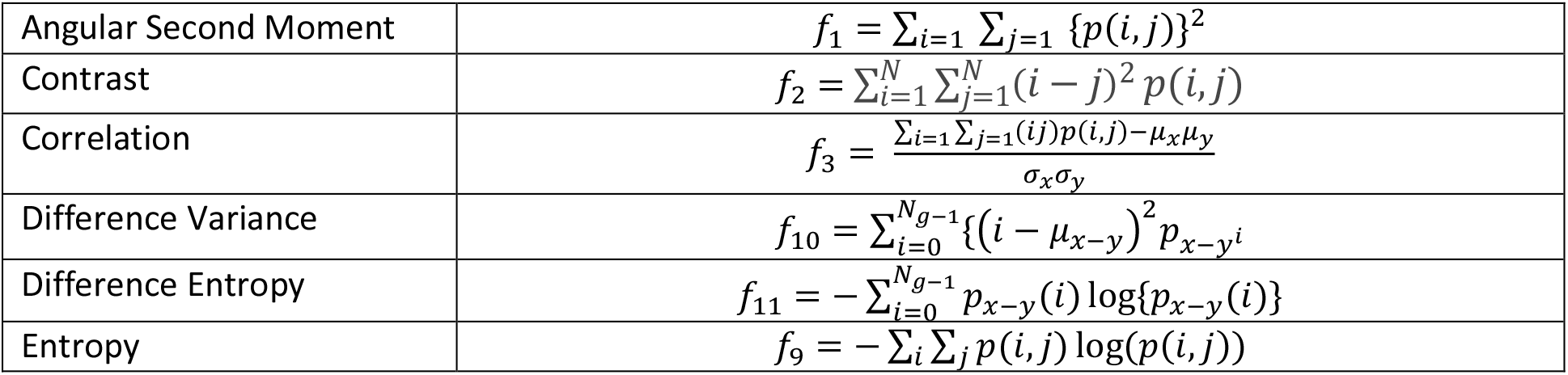
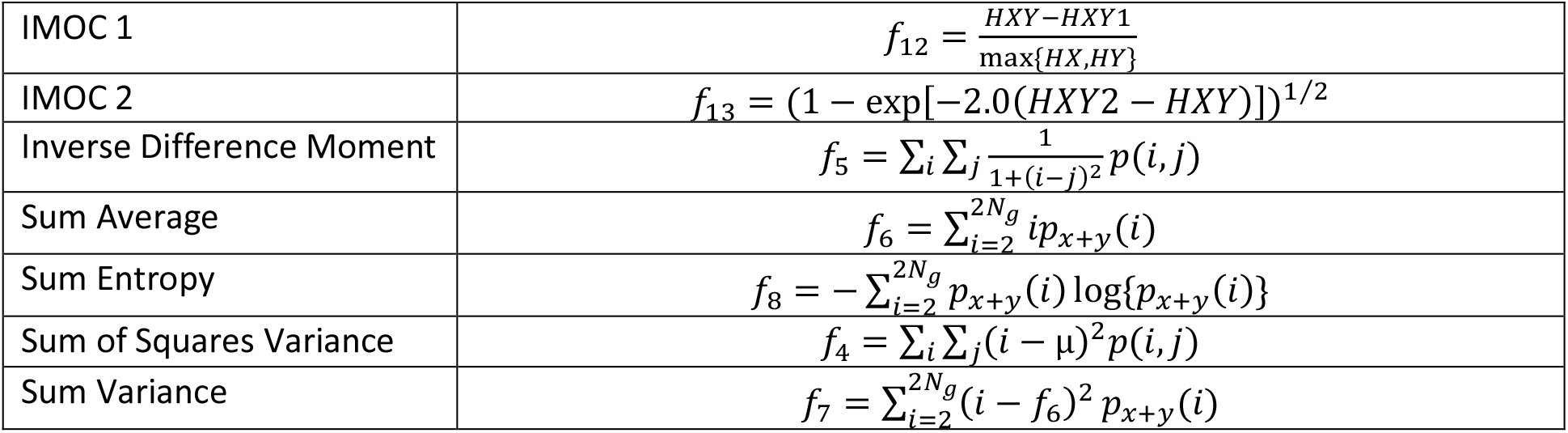
Haralick Features and their respective Equations

## Results and Discussion

Haralick et al listed 14 features for the characterization of textures, but in the case of this study 13 were evaluated, all of which were able to provide a significant difference between at least two of the categories. The three damage categories for each textural feature were then compared using statistical analysis using a consistent alpha level of .05 for all test used. A Shapiro-Wilks normality test and a Levene test for homogeneity of variances was conducted and each textural feature failed(Levene, 1960; Shapiro & Wilk, 1965). This ruled out an ANOVA test leading to a non-parametric Kruskal-Wallis test being performed to determine if there was difference between two or more of the three categories (Kruskal & Wallis, 1952). The Kruskal-Wallis test postulated that there was a difference for all 13 features. A post hoc Dunn’s test using a Bonferroni-adjusted p-value was then preformed to identify which categories were significantly different from each other(Abdi, 2007; Dunn, 1964). The features entropy, inverse difference moment, and sum average provided the most distinctions between each of the three categories and can be found in Figure 5. These three features have shown a significant difference between the smooth and cracked, smooth and wrinkled, and the cracked and wrinkled category combinations. Angular second moment was only able to distinguish the wrinkled category from the other two. The remaining nine features used in this study were only able to differentiate smooth from the cracked and wrinkled. All the features were able to differentiate the smooth category from the other two with the exception of angular second moment which was the only feature to exclusively differentiate the wrinkled from the other two categories. A summary of the statistical analysis results for all 13 features used can be found in Table 2.

**Table 2:** Statical Analysis Results for 13 Haralick features

|  | Significant Difference<br>between<br>Smooth and Cracked | Significant Difference<br>between<br>Smooth and Wrinkled | Significant Difference<br>between<br>Cracked and Wrinkled |
| --- | --- | --- | --- |
| Angular Second Moment | No | Yes | Yes |
| Contrast | Yes | Yes | No |
| Correlation | Yes | Yes | No |
| Difference Variance | Yes | Yes | No |
| Difference Entropy | Yes | Yes | No |
| Entropy | Yes | Yes | Yes |
| IMOC 1 | Yes | Yes | No |
| IMOC 2 | Yes | Yes | No |
| Inverse Difference Moment | Yes | Yes | Yes |
| Sum Average | Yes | Yes | Yes |
| Sum Entropy | Yes | Yes | No |
| Sum of Squares Variance | Yes | Yes | No |
| Sum Variance | Yes | Yes | No |

**Table 3:** Image Acquisition Parameters

|  |  |
| --- | --- |
| Camera | 24.5-megapixel Nikon z6 |
| Lens | Nikkor Z 24-70 mm lens |
| F-stop | 20 |
| Shutter Speed | 1/125 |
| ISO | 2500 |
| Lens Height | 21.4 |
| Imaging Platform | 36 cm by 30.5 cm matte black plate of metal |
| Lighting | 3x GVM-672 LED lights |
| Light Temperature | 3500 K° |
| Brightness | 50% |
| Light Temperature (measured) | 3390 K° |
| Brightness (measured) | 2890 lux |

**Figure 5:**
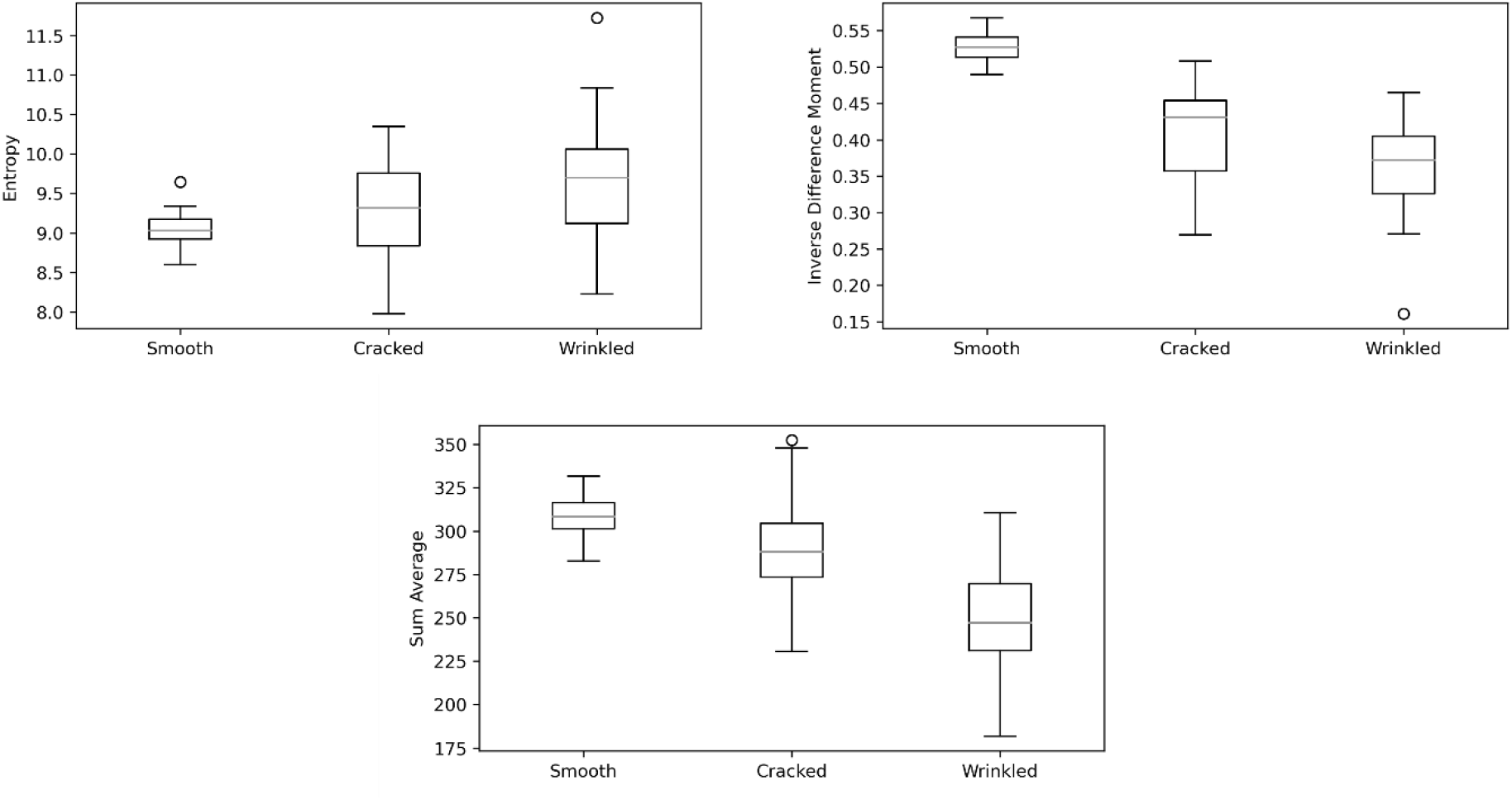
Box plot results for Entropy, Inverse Difference Moment, and Sum Average.

### Conclusion

This study has demonstrated the calculation of Haralick textural features from the surface of soybeans and their use in distinguishing and classifying three types of visual damage state in soybeans. While a majority of the Haralick features were not able to show a significant difference between all three categories, the features entropy, inverse difference moment, and sum average did show a significant difference between all three categories, while the remaining features only distinguished one category from the other two. The ability of these Haralick textural features to show a distinction between the categories may be of value to improving methods of detecting damage in grains similar to soybeans, and in further classifying them. Further research into the image analysis of other visual characteristics such as color and shape may provide more insight into further differentiating between the three categories. A combination of texture, color, and shape image analysis can improve the visual evaluation, categorization, and sorting of soybean and may show potential in adaptation to numerous other grains.

## Appendix

## Notes

### Competing Interest Statement

The authors have declared no competing interest.

